# The CoQ biosynthetic di-iron carboxylate hydroxylase COQ7 is inhibited by in vivo metalation with manganese but remains function by metalation with cobalt

**DOI:** 10.1101/2022.07.12.499729

**Authors:** Ying Wang, Siegfried Hekimi

## Abstract

COQ7 is a mitochondrial hydroxylase that catalyzes the penultimate step of the biosynthesis of coenzyme Q (CoQ; ubiquinone). CoQ is an obligate component of the mitochondrial electron transport chain and an antioxidant. CoQ deficiencies due to mutations in CoQ biosynthetic enzymes are severe genetic disorders often manifesting as mitochondrial disease syndrome. COQ7 is part of the relatively rare class of di-iron carboxylate enzymes, which carry out a wide range of reactions. In a previous study we described how COQ7 activity is inhibited in mammalian cells after treatment with iron chelating agents. Here, we report that manganese exposure of mouse cells leads to decreased COQ7 activity and resulting CoQ deficiency, which might participate in manganese toxicity. We find that the presence of cobalt can interfere with the inhibition of COQ7 by manganese. We present evidence that both manganese inhibition and cobalt interference are the result of metal exchange at the di-iron active site of COQ7. We present findings that suggest that 1) cobalt has greater affinity for the active site of COQ7 than both iron and manganese and, 2) that iron replacement by cobalt at the active site preserves catalytic activity.

## INTRODUCTION

COQ7 is the penultimate enzyme in the coenzyme Q (CoQ) biosynthetic pathway. CoQ, also known as ubiquinone, is an endogenously synthesized, highly hydrophobic, lipid-soluble molecule found in virtually all animal cells as well as in plants and microbes [1, 2]. It is a pivotal component of the mitochondrial respiratory chain, where it functions as an obligate electron carrier, enabling ATP generation from oxidative phosphorylation [3]. It has also been implicated in modulating the mitochondrial Permeability Transition Pore (mtPTP), although the exact mechanisms by which it acts on this process remains to be elucidated [4]. Outside of mitochondria, CoQ is required for the plasma membrane redox system, acting as an intermediate electron carrier accepting electrons from cytosolic NAD(PH)[3, 4]. High concentrations of CoQ are also found in lysosome and Golgi membranes [5], where its reduction/oxidation cycle is coupled with the transfer of protons across the membrane, which is essential for maintaining the transmembrane proton gradient and luminal pH [6, 7]. Furthermore, the antioxidant function of CoQ, both as a direct electron donor and via its function in the regeneration of other essential antioxidants, has been widely acknowledged [8, 9].

Loss of sufficient CoQ biosynthesis is associated with diverse severe pathologies. Primary CoQ deficiency, due to mutation(s) in any of the CoQ biosynthetic genes (COQ genes), is often a severe debilitating illness that tends to affect multiple organs and is associated with high mortality [2, 10-12]. CoQ levels can also be affected by processes that are not directly related to CoQ biosynthesis. For example, CoQ deficiency has been reported as a secondary effect in patients with mitochondrial disease syndrome [13-17]. Low CoQ levels were also observed to accompany aging and in age-related diseases, such as heart disease and Parkinson’s disease [4, 18, 19].

The CoQ biosynthetic pathway is highly conserved from yeast to humans. The final steps are carried out in the mitochondria. The currently well-accepted model of CoQ biosynthesis is that several CoQ biosynthetic pathway components form a supramolecular assembly in the inner membrane of mitochondria, called the CoQ complex, potentially facilitating substrate channeling and CoQ export out of mitochondria [16, 20, 21]. COQ7 is a part of the CoQ complex in the budding yeast *Saccharomyces cerevisiae* and is necessary for its stability [20, 22, 23]. However, in contrast to yeast, complete loss of COQ7 in animal cells results in accumulation of its substrate, demethoxyubiquinone (DMQ), which is the only CoQ intermediate detected that can accumulate because of disrupted CoQ biosynthesis [24-27].

Structurally, COQ7 has been identified to be a carboxylate-bridged di-iron enzyme [28, 29]. Di-iron carboxylate proteins are a growing family of functionally diverse proteins that characteristically contains a di-iron active center for binding and activation of dioxygen [30, 31]. Enzymes with the non-heme di-iron cofactors catalyze a variety of distinct reactions from bacteria to humans [32]. Examples include ribonucleotide reductase (*Escherichia coli* DNA biosynthesis), Δ^9-^stearoyl-acyl carrier protein desaturase (fatty acid desaturation in plants), bacterial methane monooxygenases (conversion of methane to methanol), alternative oxidase (oxidation of ubiquinol by oxygen in plants and lower eukaryotes), and human deoxyhypusine hydroxylase (regulation of cell proliferation) [33-37]. The two iron atoms in a carboxylate active site are bridged by at least one carboxylate group from either an aspartate or a glutamate residue [31]. CLK-1/COQ7 was identified as a di-iron carboxylate protein on the basis of the conservation of six amino acids (four carboxylate residues and two histidines) constituting an iron-binding motif [28, 31]. Consistent with this, we previously reported that exposure to the metal-chelating agent clioquinol inhibits CLK-1/COQ7 activity due to iron chelation [38].

A number of metals are essential for living organisms. About one-third of all proteins depend on metal ions to function properly, and are referred to as metalloproteins [39]. Furthermore, over 40% of all enzymes have been estimated to employ metals in their active sites for catalysis [40, 41]. Metalation is aided by metal transport and specialized delivery mechanisms, but the majority of metalloenzymes (70%) are thought to lack selective mechanisms for binding the appropriate metal species. Furthermore, they compete with other molecules, including other proteins, for the limited metal pools in the cell [42]. The relative affinities of proteins for divalent metal ions follow the Irving–Williams series: Mn^2+^ < Fe^2+^ < Co^2+^ < Ni^2+^ < Cu^2+^ > Zn^2+^. However, the cell can assist desired metalation by maintaining the relative availabilities of metal ions to the inverse of the Irving–Williams series with weaker binding metals such as Mn^2+^ and Fe^2+^ at greater availabilities, and the tighter binding metals like Co^2+^ at lower concentrations [43, 44]. Nevertheless, as proteins have only a limited ability to discriminate between different metal ions, there remains a risk of mis-metalation which can often lead to enzyme inactivation [43]. In fact, the ability to displace other essential metals from binding sites in proteins has been invoked as one of the key mechanisms of metal toxicity, due to the likely inhibition of protein functions.

In this present study, we report that manganese (Mn^2+^) exposure inactivates COQ7 and thus causes CoQ deficiency. This is likely caused by mis-metalation with Mn^2+^ instead of Fe^2+^ at the enzyme’s activity site. Furthermore, we present surprising findings that suggest that under the right experimental conditions Co^2+^ can replace the Fe^2+^ atoms at the active site of COQ7 and bind very strongly without compromising COQ7 activity. This could have important consequences for the utilization of other di-iron enzymes of industrial importance.

## RESULTS

### Manganese exposure results in decreased COQ7 activity

The only known function of COQ7 is hydroxylation of DMQ in CoQ biosynthesis [23-27]. We have previously shown that treatment of cells from a mouse macrophages cell line (RAW264.7) with the metal chelator clioquinol results in a decrease of COQ7 activity because of iron loss from the active site [38]. We now found that exposure of RAW264.7 cells to Mn^2+^ ions also results in CoQ loss and accumulation of DMQ, which indicates COQ7 inhibition (Figure 1A). CoQ is composed of a benzoquinone ring and a lipophilic poly-isoprenoid tail whose chain length varies according to species. Mouse cells have mostly CoQ_9_ (9 referring to the number of isoprene units in the tail) but also a small amount of CoQ_10_. We observed similar effects of Mn^2+^ exposure on CoQ_9_ and CoQ_10_ (Figure 1A).

**Figure 1.**
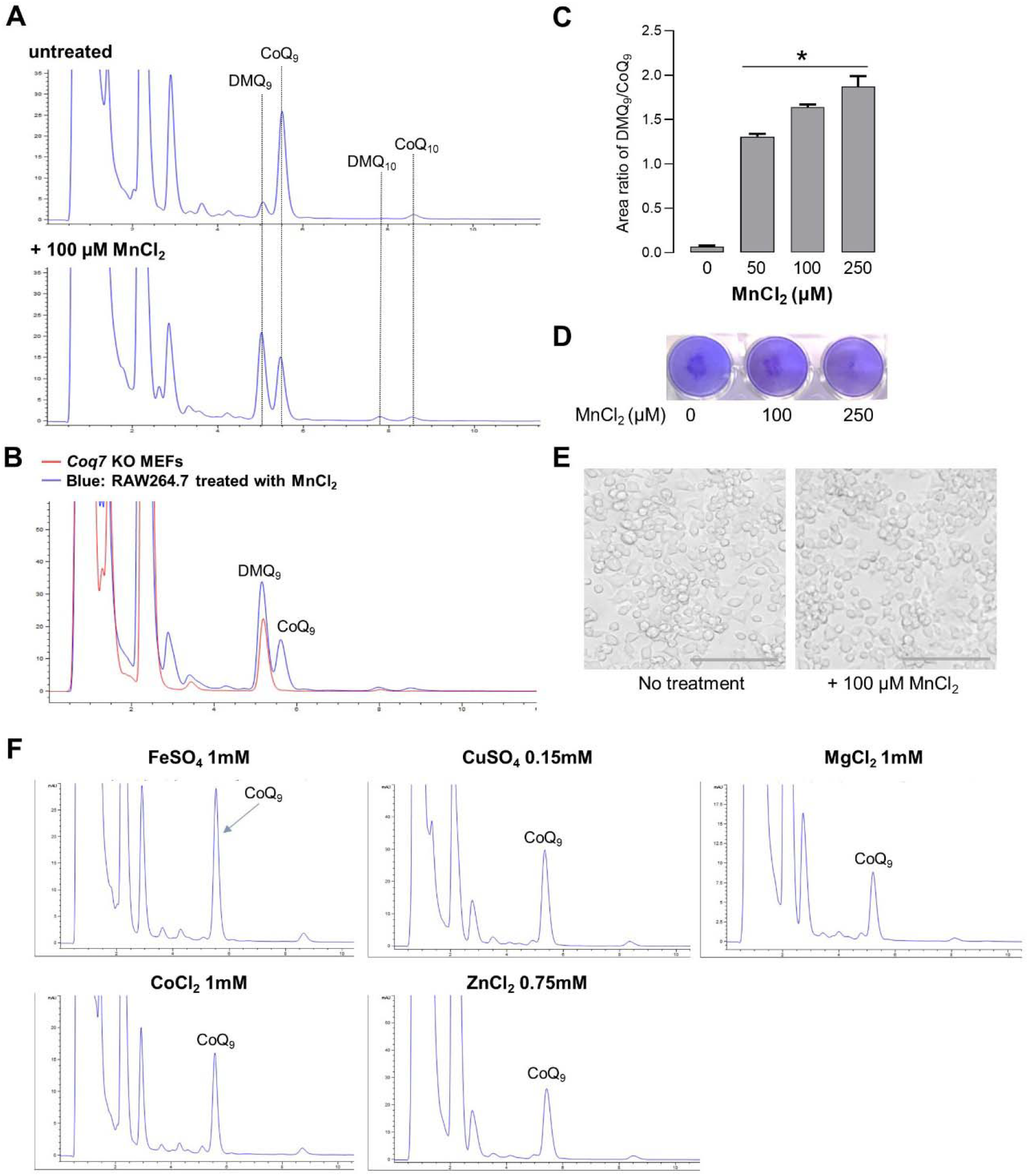
Mn^2+^ exposure results in a decrease of COQ7 activity. (**A**) High-performance liquid chromatography (HPLC) traces of CoQ extracts from MnCl_2_-treated or untreated RAW264.7 cells. (**B**) The HPLC trace of quinone extract from McCl_2_-treated RAW264.7 cells is overlaid with the extracted quinone from *Coq7* KO MEFs (24,26), confirming the identifies of the DMQ and CoQ peaks in RAW264.7 cells. (**C**) Increased DMQ_9_:CoQ_9_ ratio after MnCl_2_ treatment. Data are shown as mean ± SEM of 2 replicates. \**p*<0.05 versus untreated control (one-way ANOVA followed by Dunnett’s post hoc). MnCl_2_ was added to culture medium for 24 hours before harvest for CoQ extraction. (**D**) Crystal violet staining shows no significant loss of cell population after 24 h treatment with MnCl_2_. (**E**) Phase contrast microscopy images show no alterations in cell morphology after MnCl_2_ exposure. Scale bar = 100 µm. (**F**) Representative HPLC traces of quinone extracts after 24h treatment with different metal ions. None of the indicated treatments resulted in the appearance of DMQ, indicating a lack of effect on COQ7 activity. Of note, different amounts of cells were used for quinone extraction and therefore the sizes of CoQ_9_ peaks vary in the traces shown but do not represent effect of the ions on the total level of synthesis.

Note that a minor DMQ peak is detectable in some types of cultured cells such as RAW264.7 cells (Figure 1A). DMQ elutes just a bit earlier than CoQ, but the two compounds are readily separable into 2 distinct peaks [23, 24, 45]. Comparison with the quinone extract from mouse Coq7 KO MEFs, which only produce DMQ, allows for further confirmation of the identify of the DMQ peak [26] (Figure 1B). It is currently impossible to exactly quantify cellular DMQ concentration due to the unavailability of synthetic DMQ to use as standard. We therefore use the DMQ_9_ to CoQ_9_ ratio as the measure of COQ7 enzymatic activity. As shown in Figure 1C and S1, Mn^2+^ treatment results in a dose-dependent increase of the DMQ_9_/CoQ_9_ ratio, while no significant effect on cell viability was observed (Figure 1D and 1E). In contrast, no other metal tested, including Cu^2+^, Mg^2+^, and Co^2+^, was found to cause accumulation of DMQ (Figure 1F). Of note, the test doses for the different metals were chosen based on cytotoxicity test results and, for each metal, the maximum doses tolerated for 24h treatments were used in all relevant experiments of this study (Fig. S2).

### Co-treatment with iron prevents the inhibition of COQ7 by manganese

We speculated that inhibition of COQ7 activity by Mn^2+^ was due either to interference with Fe^2+^ uptake and/or by binding to COQ7 instead of Fe^2+^. To test this, we first determined the effect on COQ7 activity of simultaneous exposure to Mn^2+^ and Fe^2+^. We found that Fe^2+^ co-treatment significantly prevents the inhibitory effect of Mn^2+^ on COQ7 activity (Figure 2A). Mn^2+^ and Fe^2+^ might affect each other’s uptake. Possibly this could explain the inhibition of COQ7 activity by Mn^2+^ if it results in intracellular Fe^2+^ depletion. To explore this further, we pretreated the cells for 3h with Fe^2+^; then, after removal of the Fe^2+^ from the media, the cells were treated with Mn^2+^ for a total of 24h before CoQ extraction. Under these conditions, the cells should have plenty of Fe^2+^, yet this had no effect on the inhibitory effect of Mn^2+^ on COQ7 (Figure 2B). This result suggests that when co-treatment with Fe^2+^ prevents inhibition of COQ7 by Mn^2+^, it is likely mainly due to Fe^2+^ competing for the cellular uptake and binding of Mn^2+^. However, in the presence of increased intracellular concentration of Fe^2+^ as a result of Fe^2+^ pre-loading, Mn^2+^ still succeeds in inhibiting COQ7, suggesting the possibility of a preference for Mn^2+^ over Fe^2+^ for binding to the metal binding site of the enzyme.

**Figure 2.**
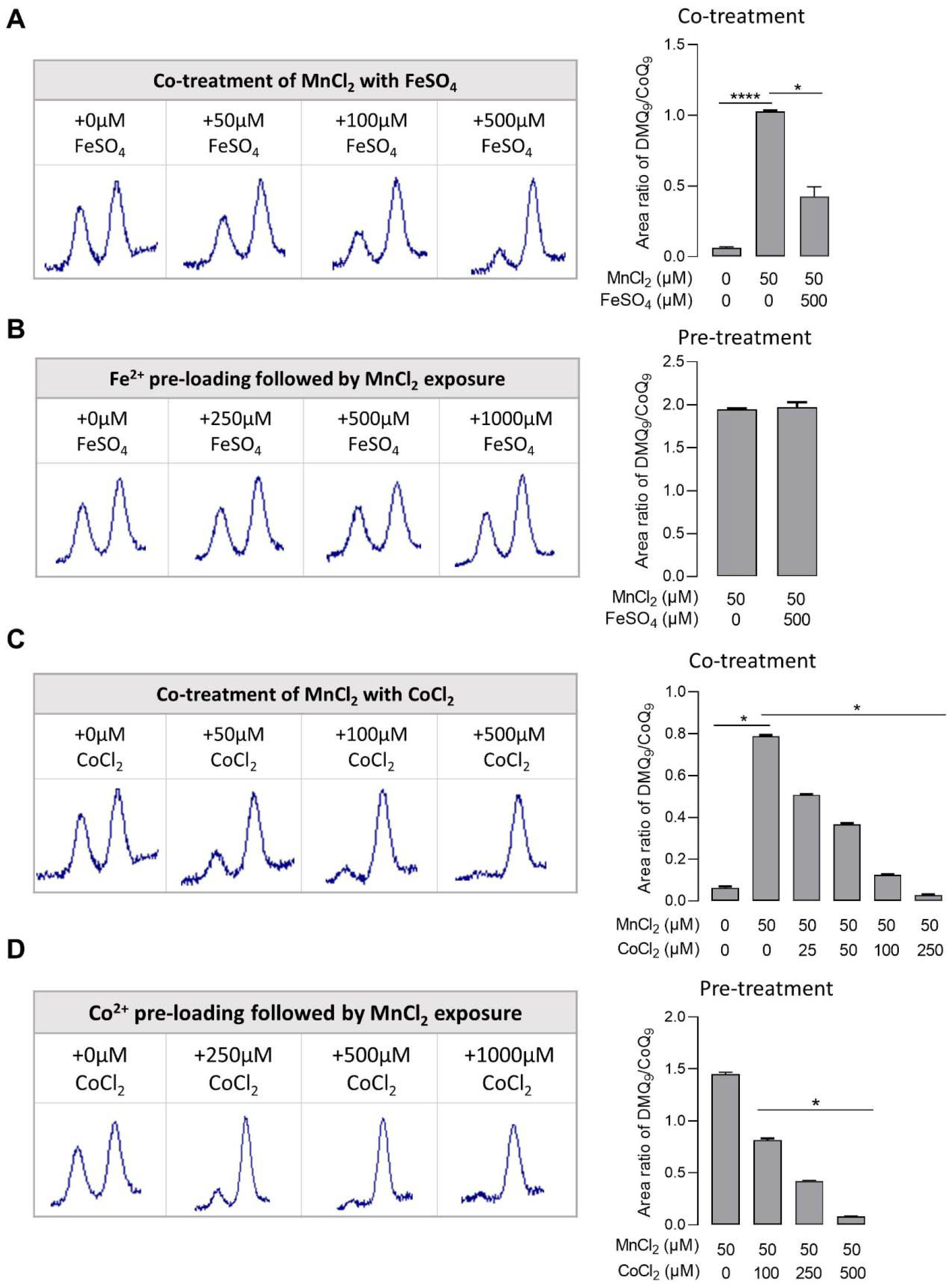
Effect of Fe^2+^ or Co^2+^ loading on the inhibition of COQ7 by Mn^2+^. Throughout the figure each of the chromatograms show, from left to right, a DMQ_9_ peak followed by a CoQ_9_ peak. Quantitative DMQ_9_:CoQ_9_ ratios are shown on the right. (**A**) Effect of FeSO_4_ co-treatment on inhibition of COQ7 activity induced by MnCl_2_ exposure. (**B**) Pre-loading cells with FeSO_4_ showed no significant effect on inhibition of COQ7 activity induced by MnCl_2_ exposure. (**C**) Effect of co-treatment with CoCl_2_ on inhibition of COQ7 activity induced by MnCl_2_ exposure. (**D**) Pre-loading cells with CoCl_2_ prevents the inhibitory effect of MnCl_2_ on COQ7 activity. The graphs on the right depict the mean ± SEM (n=3) and the statistical analysis was performed by one-way ANOVA followed by Dunnett’s post hoc test (\*\*\*\**p*< 0.0001; \**p*<0.05).

### Cobalt prevents the inhibition of COQ7 by manganese

We found that co-treatment with Co^2+^ also prevents Mn^2+^ from inhibiting COQ7, perhaps even more efficiently than Fe^2+^ (Figure 2C). As with Fe^2+^, this could be due to competition for uptake of Mn^2+^. However, surprisingly, and in contrast to what we saw with Fe^2+^, pre-loading cells with Co^2+^ prevented the inhibitory effect of Mn^2+^ on COQ7 (Figure 2D). All the other metals tested, however, including Cu^2+^, Mg^2+^, Ni^2+^ and Zn^2+^ were like Fe^2+^ unable to block the effect of Mn^2+^ (Figure 3). Treatment of human embryonic kidney (HEK-293) cells with Mn^2+^ also led to reduced activity of COQ7. The most common CoQ in human cells is CoQ_10_. As shown in Figure. S4, exposure to 0.5mM of Mn^2+^ for 24h resulted in appearance of DMQ_10_ and an increased of DMQ_10_ to CoQ_10_ ratio. However, the effects are less pronounced compared to those seen with RAW264.7 cells. Pre-loading HEK293 cells with 0.5mM of Co^2+^ also significantly attenuated the inhibition of COQ7 activity by Mn^2+^.

Like Mn^2+^ and Fe^2+^, Co^2+^ is redox-active under physiological conditions. Its properties are similar to those of Fe^2+^ and, according to the Irving-Williams series, Co^2+^ forms more stable complexes with proteins than do Fe^2+^ and Mn^2+^. Based on these facts, we postulate that Co^2+^ can substitute for Fe^2+^ at the active site of COQ7 without compromising its activity. Furthermore, as Co^2+^ has a higher binding constant for proteins than Mn^2+^, Mn^2+^ cannot displace Co^2+^ at the active site of COQ7, which is why Co^2+^ pre-loading can prevent Mn^2+^ inhibition of COQ7.

**Figure 3.**
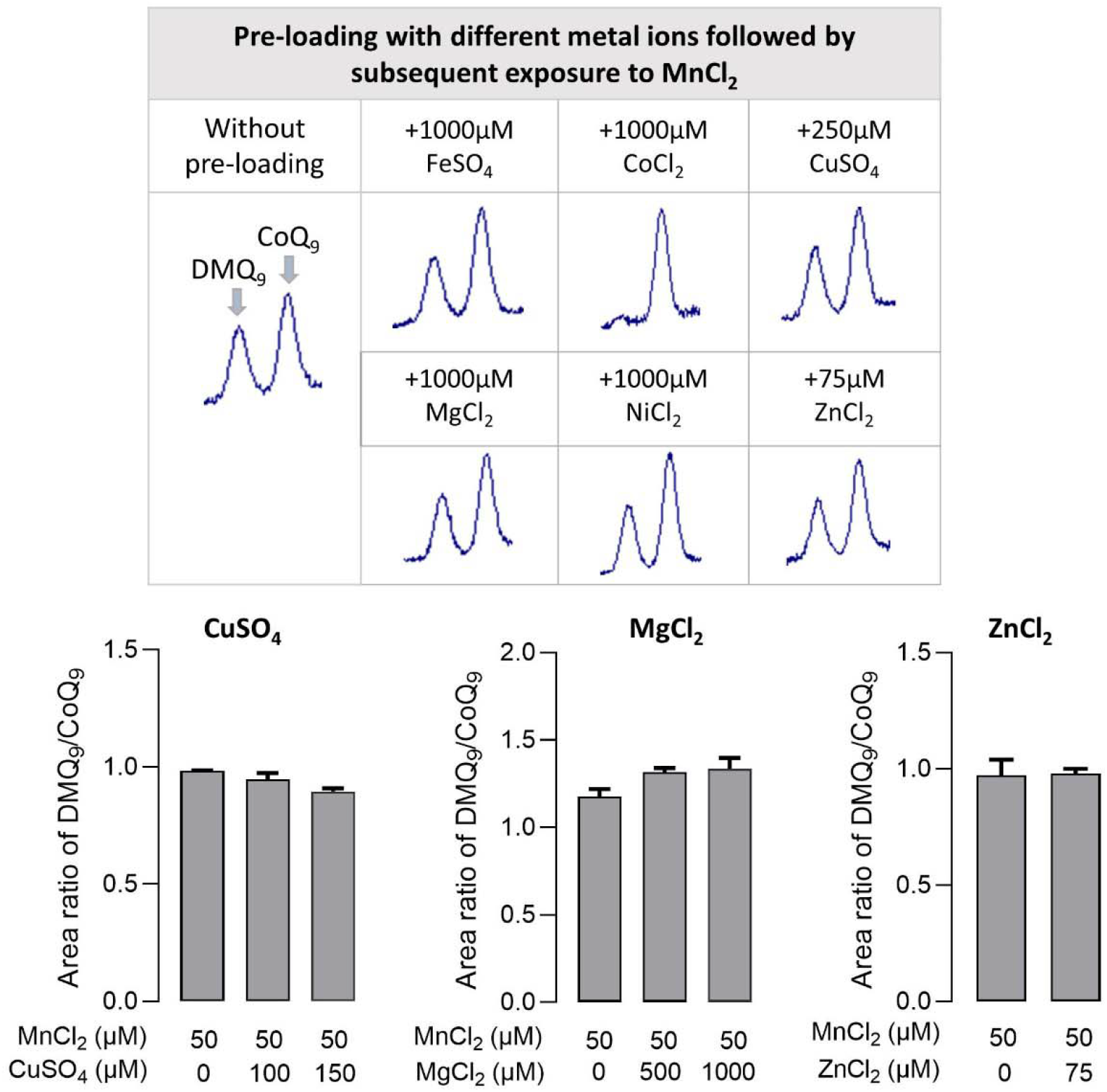
Pre-loading with Cu^2+^, Mg^2+^ or Zn^2+^ shows no effect on the inhibition of COQ7 activity by Mn^2+^. RAW 264.7 cells were treated with the indicated metal icons for 3h before the metal ions were removed from the media and the cells were cultured for an additional 24h with MnCl_2_ (50 µM). Representative HPLC traces of DMQ_9_ and CoQ_9_ peaks and the peak area ratio of DMQ_9_ to CoQ_9_ at the end of treatments are shown. Error bar is SEM, n = 3.

### Effects of cobalt and iron on the recovery of COQ7 activity after removal of manganese

We found that COQ7 activity slowly recovers after removal of Mn^2+^ from the medium (Figure 4A). Addition of Fe^2+^ in the medium after Mn^2+^ washout significantly accelerates the recovery of COQ7 activity (Figure 4B), consistent with our hypothesis that inhibition of COQ7 activity by Mn^2+^ is by substituting for Fe at the di-iron active site of the enzyme. Co^2+^ addition shows the same effect (Figure 4B), which is again consistent with the notion that Co^2+^ is as functional as Fe^2+^ at the active site of COQ7.

**Figure 4.**
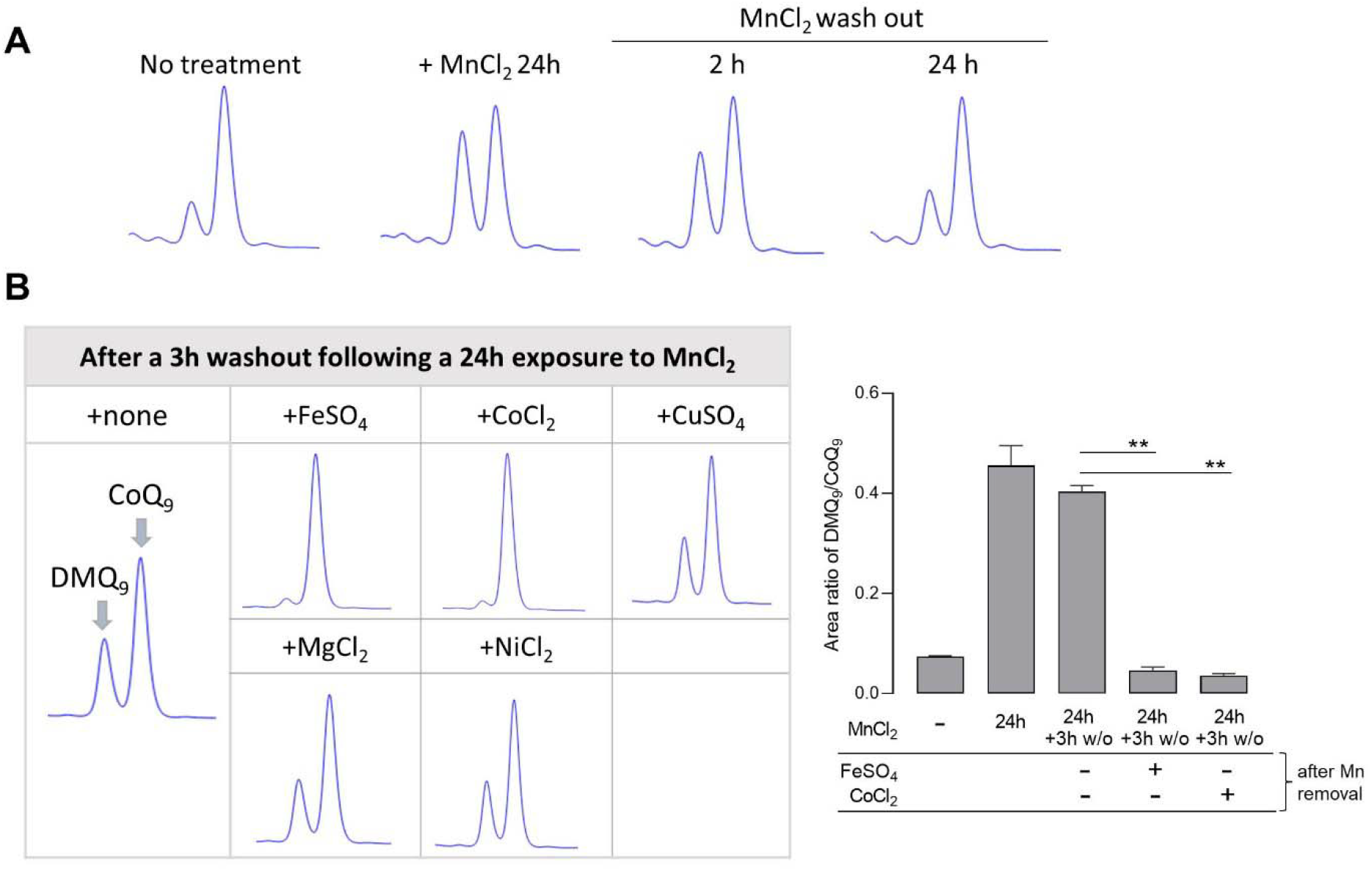
Reversal of COQ7 activity following the removal of Mn^2+^ from the medium. (**A**) HPLC traces of DMQ_9_ and CoQ_9_ peaks showing a partial recovery of COQ7 activity 24h after washout of MnCl_2_ from the medium. (**B**) Supplementing FeSO_4_ or CoCl_2_ in the medium for 3h after the washout of Mn^2+^ resulted in complete recovery of COQ7 activity, while treatments with other indicated metal icons (Cu^2+^, Mg^2+^ or Ni^2+^) showed no difference compared to no-treatment control. MnCl_2_: 50 µM. Post-Mn^2+^ exposure with other metal salts: 500 µM. Error bar is SEM, n = 3. \*\**p*< 0.01 (one-way ANOVA with Dunnett’s multiple comparison test).

Unfortunately, we are unable to directly demonstrate the presence of cobalt in active COQ7 protein. First, it is difficult to isolate metalated proteins to identify endogenous metal cofactors because metal ions tend to be lost or replaced during the process. Furthermore, COQ7 is associated with the mitochondrial inner membrane which makes it more challenging to purify. Finally, no in vitro activity assay exists for COQ7. Thus, we could not hope to reconstitute transgenic COQ7 with metals in vitro and determine whether it is active.

### Mn^2+^ exposure increases COQ7 protein level

We used Western blot analysis to determine whether the inhibition of COQ7 activity by Mn^2+^ is associated with a change of COQ7 expression level. The COQ7 level was found to be higher in Mn^2+^-treated cells and the change could not be prevented by co-treatment with Co^2+^, despite its ability to maintain COQ7 functionality (Figure 5). The effect of Mn^2+^ is specific for COQ7 as the level of PDSS2 was not affected (Figure 5). PDSS2 is an another COQ-biosynthetic enzyme, which catalyzes the assembly of the polyisoprenoid side chain. Co^2+^ treatment at a relatively high dose (250 µM), also leads to elevation of COQ7 concentration. We speculate that the increase of COQ7 level after Mn^2+^ or Co^2+^ exposure could be due to increased COQ7 protein stability because of Mn or Co at the enzyme’s active center.

**Figure 5.**
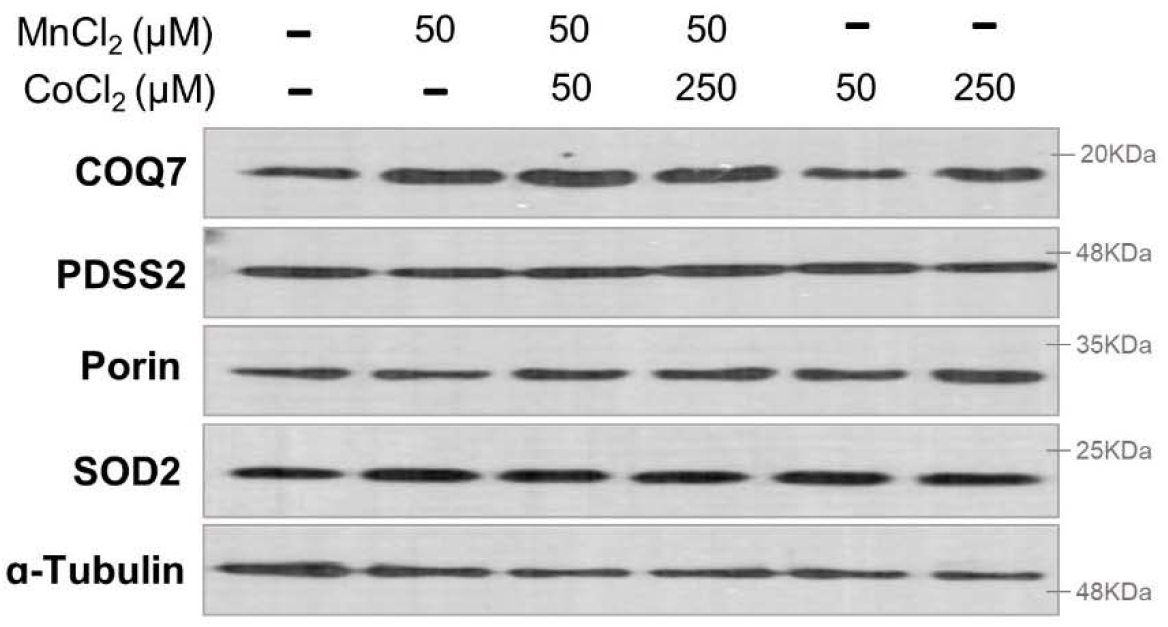
Western blot analysis of COQ7. Western blots showing increased COQ7 protein level after 24h MnCl_2_ treatment. COQ7 level was also elevated by CoCl_2_ treatment at 250 µM. Simultaneous exposure to CoCl_2_ showed no effect on the elevation of COQ7 level by MnCl_2_. The original scans are presented in Supplemental Fig. S3.

## DISCUSSION

The CoQ biosynthetic enzyme COQ7 requires Fe in its active center to catalyze DMQ hydroxylation [28, 29, 38, 46]. Our study demonstrates that Mn^2+^ exposure leads to loss of COQ7 activity and CoQ deficiency. Fe and Mn are two of the most important biologically relevant metals. For humans, along with Co, Cu, and Zn, they are the 5 essential trace elements of the 10 first-row transition metals. Both are relatively weak binders to metalloproteins as predicted by the Irving Williams series [44]. However, their intracellular concentrations are relatively high, and Fe and Mn are used by about 8% and 6% of all enzymes, respectively [44]. Divalent ions of Mn and of Fe are similar in chemistry, and both tend to coordinate with six ligands and from octahedral complexes [47]. This makes it difficult for proteins to discriminate between Fe^2+^ and Mn^2+^ on the basis of structure alone [47]. Therefore, when local availability of Mn^2+^ is above the physiological level, there is a high likelihood that Mn^2+^ may outcompete Fe^2+^ for metal-binding sites of proteins, even of those that cannot function with Mn^2+^ [47]. As an example, Mn^2+^ overload was shown to inhibit aconitase’s catalytic function, which was postulated to be at least in part because of competition with Fe^2+^ for binding to the iron-sulfur cluster that directly reacts with the enzyme substrate citrate [48, 49]. We thus postulate that the inhibitory effect of manganese on COQ7 is mainly by replacing the iron atoms at the enzymatic active site of COQ7.

Alternatively, inhibition of COQ7 by Mn^2+^ exposure could be due to competition with Fe^2+^ for cellular and mitochondrial uptake. The exact absorption and transport mechanisms for different metals are not fully understood. However, shared transport and trafficking mechanisms have been reported. Mn is thought to share transporters with Fe, and specific transporters and regulatory proteins have also been identified [50]. For example, DMT1 is a transmembrane glycoprotein which mediates the proton-coupled transport of various divalent metals, including Fe^2+^, Mn^2+^, and others, across multiple organellar membranes, including the plasma membrane, and the mitochondrial and Golgi membranes [51, 52]. The voltage gated ZIP8 channels also pump various metals including Co^2+^, Fe^2+^, Mn^2+^ across membranes [51, 52]. Both Fe and Mn have been shown to use the calcium uniporter to cross the inner membrane into the matrix [53-55]. COQ7 is a part of the CoQ biosynthetic complex on the matrix face of the inner mitochondrial membrane [16, 20, 56, 57]. It’s not known where and how the metalation of COQ7 takes place. However, mitochondrial metalloproteins are generally believed to acquire their metals after import into the organelle [43]. It is possible that increased Mn availability interferes with the Fe uptake capacity of mitochondria leading to insufficient Fe levels in the mitochondria for binding to the active site of COQ7. However, the observation that COQ7 is still affected in the cells loaded with excess of Fe^2+^ before Mn^2+^ exposure weakens the possibility that competition for uptake is the main mode of action for COQ7 inhibition by Mn^2+^ exposure. Rather, it suggests that COQ7 may preferentially binds to Mn^2+^ over Fe^2+^ when Mn^2+^ is sufficiently available The observation that after stopping Mn^2+^ treatment, supplementing culture medium with Fe^2+^ speeds up the recovery of COQ7 activity is also in line with the metal replacement hypothesis. Toxic effects of excess Mn^2+^ are well known, especially in the central nervous system [58]. Our study suggests that Mn^2+^ toxicity could lead to CoQ deficiency.

Surprisingly, we found that in both co- and pre-treatment experiments, Co^2+^ significantly prevents the inhibition of COQ7 through Mn^2+^ exposure. Furthermore, like Fe^2+^, addition of Co^2+^ accelerates the reversal of COQ7 activity after removal of Mn^2+^ from the treatment medium. Co^2+^ has similar chemical properties as those metals in the same group on the periodic table, including Mn^2+^ and Fe^2+^. However, it occurs much less frequently in metalloproteins, likely due to its low abundance in nature [59]. Its best-known function so far is as a metal component of vitamin B_12_. We postulate that the way Co^2+^ prevents Mn^2+^ from inhibiting COQ7 and accelerates the rate of COQ7 recovery after interruption of Mn^2+^ treatment is that Co^2+^ can functionally replace the lower order metals (Fe^2+^ and Mn^2+^) at the catalytic site of COQ7, at least under the condition of an excess of Co^2+^. More specifically, as Co^2+^ lies towards the tighter binding end of the Irving–Williams series comparing to Mn^2+^, the binding of Co^2+^ to the di-iron cluster of COQ7 would render subsequent Mn^2+^ treatment less effective at inhibiting COQ7. This is because Mn^2+^ would not be able to compete out Co^2+^ due to its relatively lower affinity for binding. After interruption of Mn^2+^ treatment, the presence of Co^2+^ would allow Mn replacement by Co, leading to rapid restoration of COQ7 activity. Lastly, as 3-hour pre-loading cells with Co^2+^ is sufficient to counter the inhibitory effect of Mn^2+^ on COQ7, it is likely that Co^2+^ can in fact replace Fe^2+^ directly in the assembled COQ7 enzyme and prevent Mn^2+^ binding.

Previous studies have pointed out that protein binding sites could become mis-metalated with Co^2+^, and that this could be a major cause of Co toxicity [59]. For example, Co^2+^ was shown to directly compete with Fe^2+^ during synthesis of Fe–S clusters in *Escherichia col*i [60]. In mammalian cells, it was shown that Fe^2+^ in the Fe-binding centre of HIF-prolyl hydroxylases is replaceable with Co^2+^, resulting, in this case, in the inactivation of the hydroxylation activity [61]. On the other hand, the Fe enzyme ribulose-5-phosphate 3-epimerase (Rpe) in *Escherichia col*i was shown to be activable also by Co^2+^, Mn^2+^, and Zn^2+^, and the Mn^2+^, Fe^2+^, and Co^2+^ metalloforms of Rpe provide roughly comparable activity [62].

It is worth noting that under normal conditions, exposure of cells to CoCl_2_ does not result in significant fluorescence quenching in mitochondria in a competitive binding calcein-AM assay. Calcein-AM is a fluorescent dye whose binding to divalent cations, including Fe^2+^ and Co^2+^, results in quenching of its fluorescence. Thus, the quenching of the cytosolic signal without affecting the mitochondrial fluorescence after loading with CoCl_2_ is interpreted to indicate that Co^2+^ cannot enter mitochondria, at least not in substantial amounts [63]. However, Co^2+^ entry into mitochondria at a level far below the detection limit of the calcein-AM assay might be sufficient if the affinity of Co^2+^ for COQ7 is sufficiently high. In fact, the metal transporter DMT1 has been reported to be present in the outer membrane of mitochondria. This transporter is able to transport Co^2+^ as well as other metals. And its activity should be facilitated by high intracellular concentrations of Co^2+^ [51].

Metal ions play key structural and functional roles for nearly half of all known proteins. A better understanding of how metalation is controlled and how it modulates the functions of proteins is of importance and of growing interest. COQ7 presents a unique opportunity to look at in vivo activity with different metals because of the ease with which the reaction products produced in vivo can be monitored. Its sensitivity to metals appears to be more prevalent in the RAW264.7 macrophage line probably because of a greater propensity of macrophages to take up and accumulate metals. Nevertheless, we believe that our study is the first to describe metal swapping in a di-iron carboxylate enzyme, which are enzymes that are of commercial and ecological interest [64-66]. It warrants further studies and research into the metalation and mismetalation of di-iron carboxylate proteins.

## Material and Methods

### Cell culture and reagents

RAW264.7 purchased from the ATCC were cultured in high glucose DMEM (Dulbecco’s modified Eagle’s medium) supplemented with 10% fetal bovine serum and 1% antibiotic/antimycotic mix (Wisent, Inc) at 37°C in a humidified atmosphere of 95% air and 5% CO_2_. For single reagent treatments or co-treatment experiments, reagents were added once to the culture in 6 well plates and cells were harvested after 24h of incubation. In pre-treatment experiments, cells were exposed to different metal ions for 3h, washed with phosphate buffered saline (PBS) and then treated with MnCl_2_ for 24 hours. To test the effects of different metal ions on recovery of COQ7 activity after MnCl_2_ removal, MnCl_2_ - supplemented medium was removed after 24h of incubation. Following washes in PBS, fresh media containing different metal salts were added and CoQ was extracted at 3h after the Mn^2+^ washout. Fixed cell images were taken on an Olympus BX63 microscope using an EXi BlueTM camera and the CellSens Dimension software. All chemicals were obtained from Sigma-Aldrich.

### Extraction and high-performance liquid chromatography (HPLC) determination of CoQ

CoQ extraction and quantitation using HPLC were performed as described previously. Briefly, cell lysates were prepared in a radioimmunoprecipitation buffer (20mM Tris-HCl, pH 7.5, 1% NP-40, 0.5% deoxycholate, 10 mM EDTA, 150 mM NaCl) and CoQ were extracted with ethanol and hexane (v/v 2/5). An Agilent 1260 Infinity LC system equipped with a quaternary pump (G7111A) and a variable wavelength detector (G7114A) was used. Chromatography was carried out on a reverse-phase C18 column (2.1 × 50 mm, 1.8 µm, Agilent) with 70% methanol and 30% ethanol as the mobile phase at a flow rate of 0.3⍰mL/min. The detector was set at 275 nm. The CoQ_9_ peak was identified using pure CoQ_9_. The DMQ_9_ peak was identified by comparing to quinone extraction from Coq7 knockout mouse embryonic fibroblasts (MEFs) [24, 26].

### Western blot

Protein samples were prepared in RIPA buffer (#9806, Cell Signaling Technology), and 50 µg of lysates were subjected to 12% SDS-PAGE and visualized by using antibodies against COQ7 (15083-1-AP, ProteinTech), PDSS2 (13544-1-AP, ProteinTech), SOD1 (ab16831, Abcam), Porin/VDAC (#4661, Cell Signaling Technology), or α-Tubulin (T9026, Sigma-Aldrich). Detection was performed with Western Lightning® Plus-ECL substrate (NEL103001EA, FroggaBio) and exposed to x-ray film (sc-201697, Santa Cruz Biotechnology).

### Statistical analysis

Statistical analysis and graphical presentation were carried out using GraphPad Prism 9.2 software. A p-value of < 0.05 was considered significant for all tests.

## Author Contributions

SH and YW designed the study and wrote the manuscript together. YW performed the experiments and data analyses.

