## Supplemental Figures for "The CoQ biosynthetic di-iron carboxylate hydroxylase COQ7 is inhibited by in vivo metalation with manganese but remains function by metalation with cobalt"

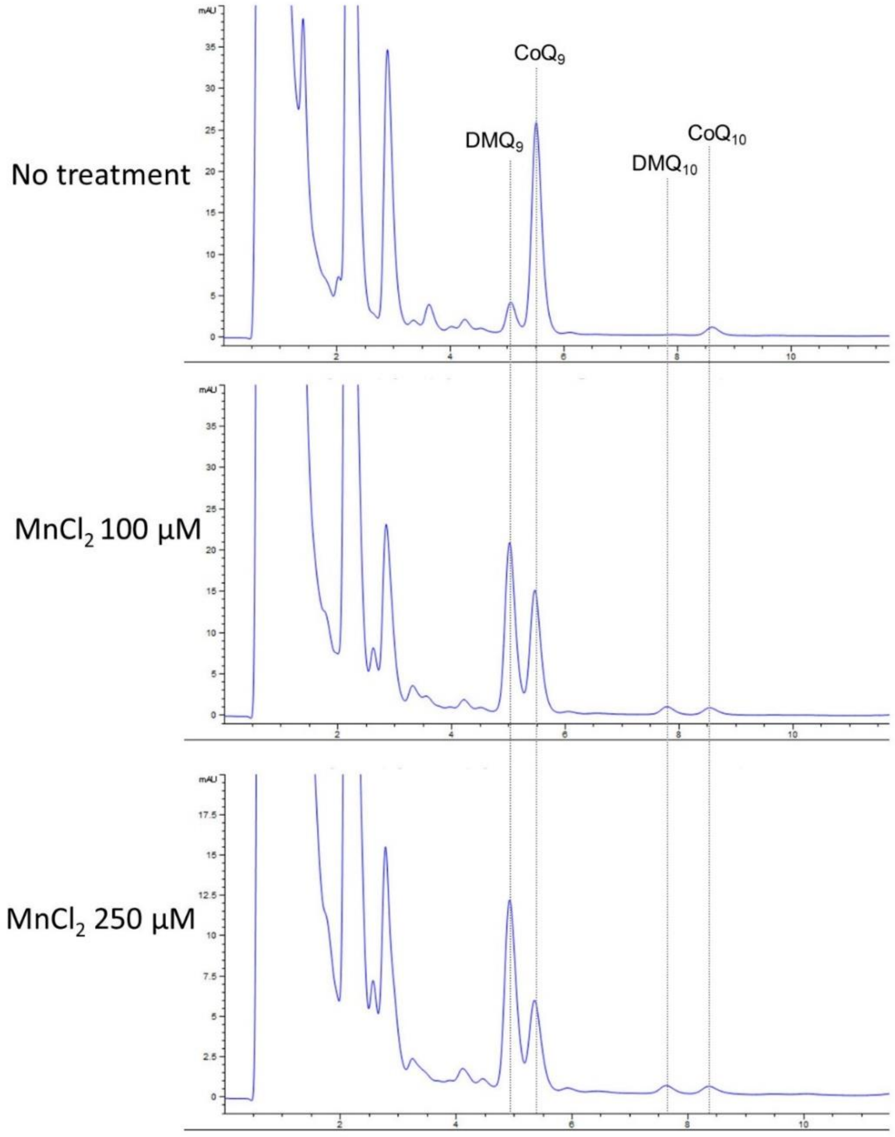


**Figure S1. $DMQ_9$ appearance and loss of $CoQ_9$ after 24 h treatment with $MnCl_2$.** HPLC traces of quinone extracts from RAW 264.7 cells are shown.

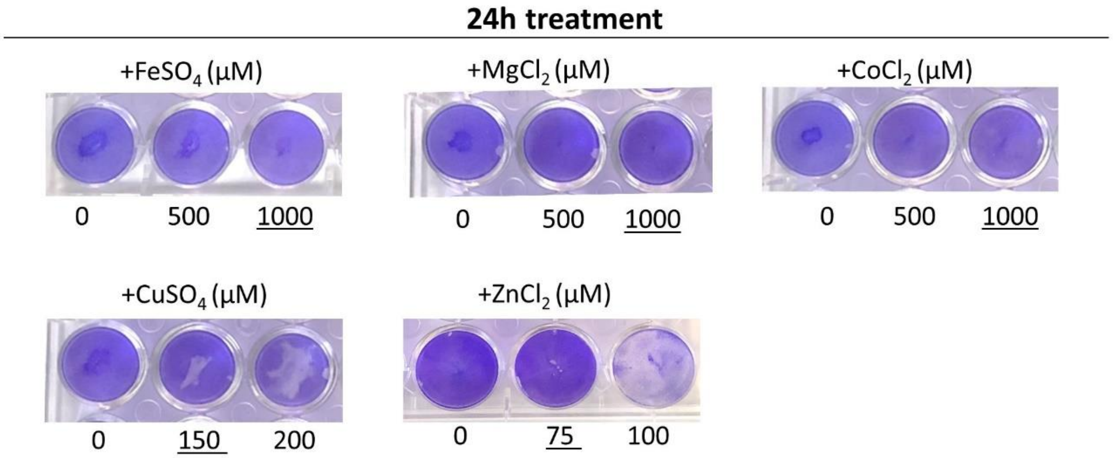


**Figure S2. Crystal violent staining for cell viability after 24h exposure to different metal ions.** For each metal ion, the maximum non-toxic dose (underlined) was chosen to use for testing its effect on the quinone content (Fig. 1F).

Antibody: COQ7, Porin

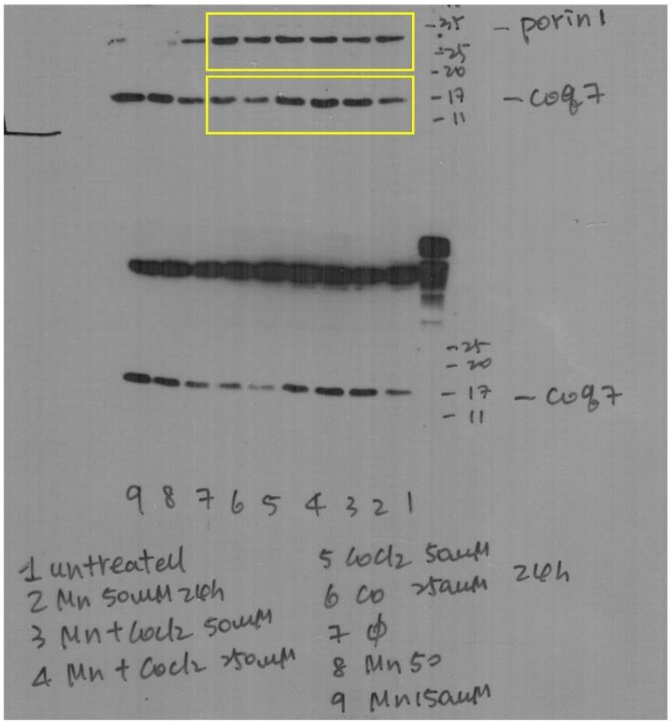


Antibody: α-Tubulin

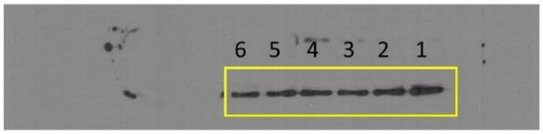


Antibody: PDSS2

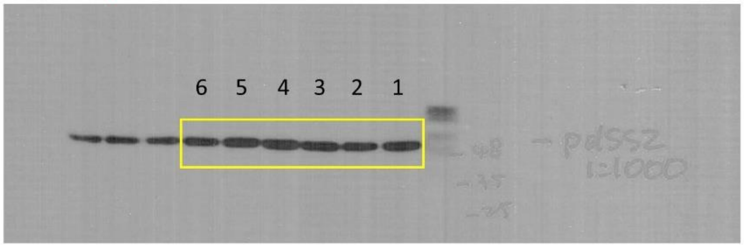


Antibody: SOD2

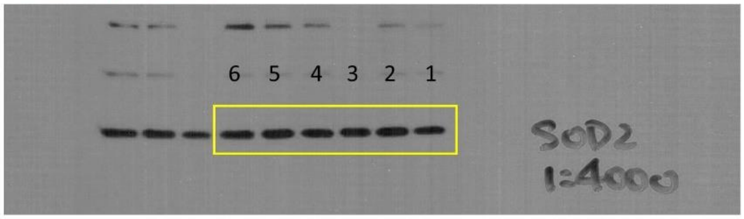


**Figure S3. Full scans of western blots presented in Figure 5.**

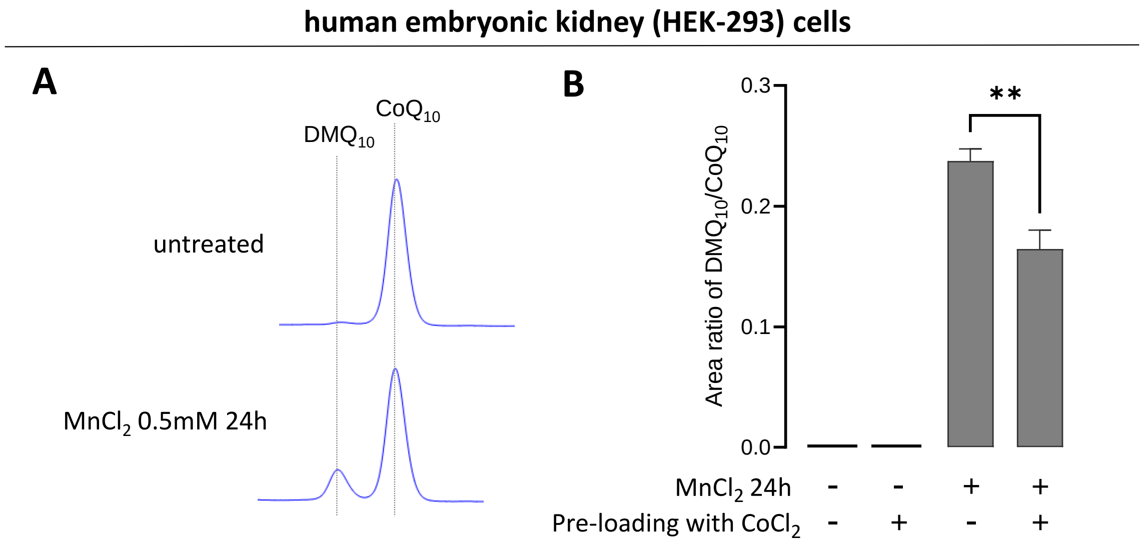


**Figure S4. $DMQ_{10}$ appearance in human embryonic kidney (HEK-293) cells after exposure to $Mn^{2+}$.** HEK-293 cells were treated with 0.5mM $MnCl_2$ for 24h before CoQ extraction. 0.5mM dose was selected because it was the highest non-toxic dose for 24h exposure of HEK-293 cells to $MnCl_2$. HPLC traces of quinone extracts are shown in A. In B, to test the effect of $Co^{2+}$ pre-loading, HEK-293 cells were treated with $CoCl_2$ for 3h followed by removal of $Co^{2+}$ from the medium, washing, and treatment with $Mn^{2+}$ for 24h.
